# Investigation of the *Cyanothece* nitrogenase cluster in *Synechocystis*: a blueprint for engineering nitrogen fixing photoautotrophs

**DOI:** 10.1101/2025.01.20.633913

**Authors:** Deng Liu, Anindita Bandyopadhyay, Michelle Liberton, Himadri B. Pakrasi, Maitrayee Bhattacharyya-Pakrasi

**Affiliations:** Department of Biology, Washington University, St. Louis, MO 63130, USA

**Keywords:** Nitrogen fixation, cyanobacteria, reconstruction, essential *nif* genes

## Abstract

The nitrogenase gene cluster of unicellular diazotrophic cyanobacteria such as *Cyanothece* is frequently selected by nature for nitrogen fixing partnerships with eukaryotic phototrophs. The essential cluster components that confer an advantage in such partnerships remain under explored. To use this cluster for the development of synthetic, phototrophic nitrogen-fixing systems, a thorough and systematic analysis of its constituent genes is necessary. An initial effort to assess the possibility of engineering this cluster into non-diazotrophic phototrophs led to the generation of a *Synechocystis* 6803 strain with significant nitrogenase activity. In the current study, a refactoring approach was taken to determine the dispensability of the non-structural genes in the cluster and to define a minimal gene set for constructing a functional nitrogenase for phototrophs. Using a bottom-up strategy, the *nif* genes from *Cyanothece* 51142 were re-organized to form new operons. The genes were then seamlessly removed to determine their essentiality in the nitrogen fixation process. We demonstrate that besides the structural genes *nifHDK*, *nifBSUENPVZTXW* as well as *hesAB* are important for optimal nitrogenase function in a phototroph. We also show that optimal expression of these genes is crucial for efficient nitrogenase activity. Our findings provide a solid foundation for generating synthetic systems that will facilitate solar powered conversion of atmospheric nitrogen into nitrogen rich compounds, a stride towards a greener world.

**IMPORTANCE:** Integrating nitrogen fixation genes into various photosynthetic organisms is an exciting strategy for converting atmospheric nitrogen into nitrogen rich products in a green and energy efficient way. In order to facilitate this process, it is essential that we understand the fundamentals of the functioning of a prokaryotic nitrogen fixing machinery in a non-diazotrophic, photoautotrophic cell. This study examines a nitrogenase gene cluster that has been naturally selected on multiple occasions for a nitrogen fixing partnership by eukaryotic photoautotrophs and provides a basic blueprint for designing a photosynthetic organism with nitrogen fixing ability.

---

Multiple examples of natural endosymbiotic events suggest a propensity for the nitrogenase (*nif*) gene cluster of unicellular cyanobacteria of the order Chroococcales to be selected for photosynthetic-diazotrophic partnerships. Of these, the *Cyanothece nif* cluster has been referred to as a ‘privileged’ partner, forming associations with diatoms such as *Rhopalodia*, *Epithemia* and *Climacodium* as well as the coccolithophore *Braarudosphaera bigelowii* (1–5). These naturally occurring instances suggest that the *Cyanothece nif* cluster harbors features that are conducive for nitrogenase to function in a photoautotroph and demands a thorough investigation as we intensify our efforts towards developing synthetic nitrogen and carbon fixing systems. In a first of its kind effort in this direction, the unicellular non-diazotrophic cyanobacterium *Synechocystis* sp. PCC 6803 (hereafter *Synechocystis* 6803) was engineered to fix nitrogen by introduction of the nitrogenase gene cluster from *Cyanothece* sp. ATCC 51142 (6). The resulting strain exhibited significant rates of nitrogen fixation, affirming the potential of this cluster to generate functional nitrogen fixing machinery in photoautotrophs and initiating our next round of efforts into its more in-depth analysis.

The commonly occurring nitrogenase enzyme is composed of two metalloproteins: the MoFe protein (dinitrogenase, NifDK) and the Fe protein (dinitrogenase reductase, NifH) (7). Besides these structural subunits of the enzyme, three iron-sulfur clusters (F cluster, P cluster, and M cluster), are required to form a mature nitrogenase enzyme complex (8, 9). The process of assembly and insertion of these three iron-sulfur clusters into the nitrogenase enzyme is complicated, involving multiple steps (10). The genetic elements required for the nitrogenase enzyme to mature vary in different diazotrophic species (11, 12). Previous efforts to engineer *E. coli* with nitrogen fixation experimented with various combinations of *nif* genes as a strategy to optimize the activity. The study identified 9 genes from the *nif* cluster of *Paenibacillus* sp. WLY78, including *nifHDK and nifBENXhesAnifV*, as essential (13). A different study determined 10 genes (*anfHDGK* and *nifJUSVFB*) from the *nif* clusters of *Azotobacter vinelandii* and *Klebsiella oxytoca* to be essential for nitrogenase activity (14). On the contrary, a recent report by Yang *et al.* identified 13 genes including *nifHDK* and *nifENBYUSVFMJ* as the essential components of the nitrogenase cluster of *Klebsiella oxytoca* (15). Nitrogenase activity could be further improved 10 fold using a refactoring approach and expressing 18 *nif* genes in various combinations along with additional genes and by altering the expression ratios of genes (16).

In our earlier work, we demonstrated the feasibility of expressing the native 24 gene *nif* cluster from *Cyanothece* 51142 in *Synechocystis* 6803. The additional 7 genes in the cluster were not included in this study (6). Further, except for the structural genes *nifHDK*, the essentiality of the other genes in the cluster in nitrogen fixation was not determined in this photosynthetic chassis. In the current study, we reconstructed *nif* gene operons using a bottom-up strategy in *Synechocystis* 6803. Promoters, ribosome binding sites (RBS), and transcriptional terminators previously identified (17) were used to control the expression of the *nif* genes. We found that 16 genes organized into 5 engineered operons formed a functional nitrogenase enzyme complex with remarkable activity. More importantly, using the CRISPR/cpf1 strategy (18), we strategically removed each gene except *nifHDK* from the reconstructed *nif* operons to determine their essentiality. Our study showed that deletion of any of these genes leads to a decrease in nitrogenase activity, thus establishing the minimal gene set required for optimal nitrogenase function in a non-diazotrophic photoautotroph.

## RESULTS

### Expression levels of structural genes affect nitrogenase activity

In the *nif* cluster of *Cyanothece* 51142, the transcriptional start sites and operons are not well defined. The only well characterized operon in this cluster is the *nifHDK* operon, the expression of which is driven by the sequence before the *nifH* gene (19). In order to test the effect of the expression strength of these genes on nitrogenase activity, we cloned the sequence between *nifU* and *nifH* into cassettes containing different promoters (Fig. 1A).

**FIG 1.**
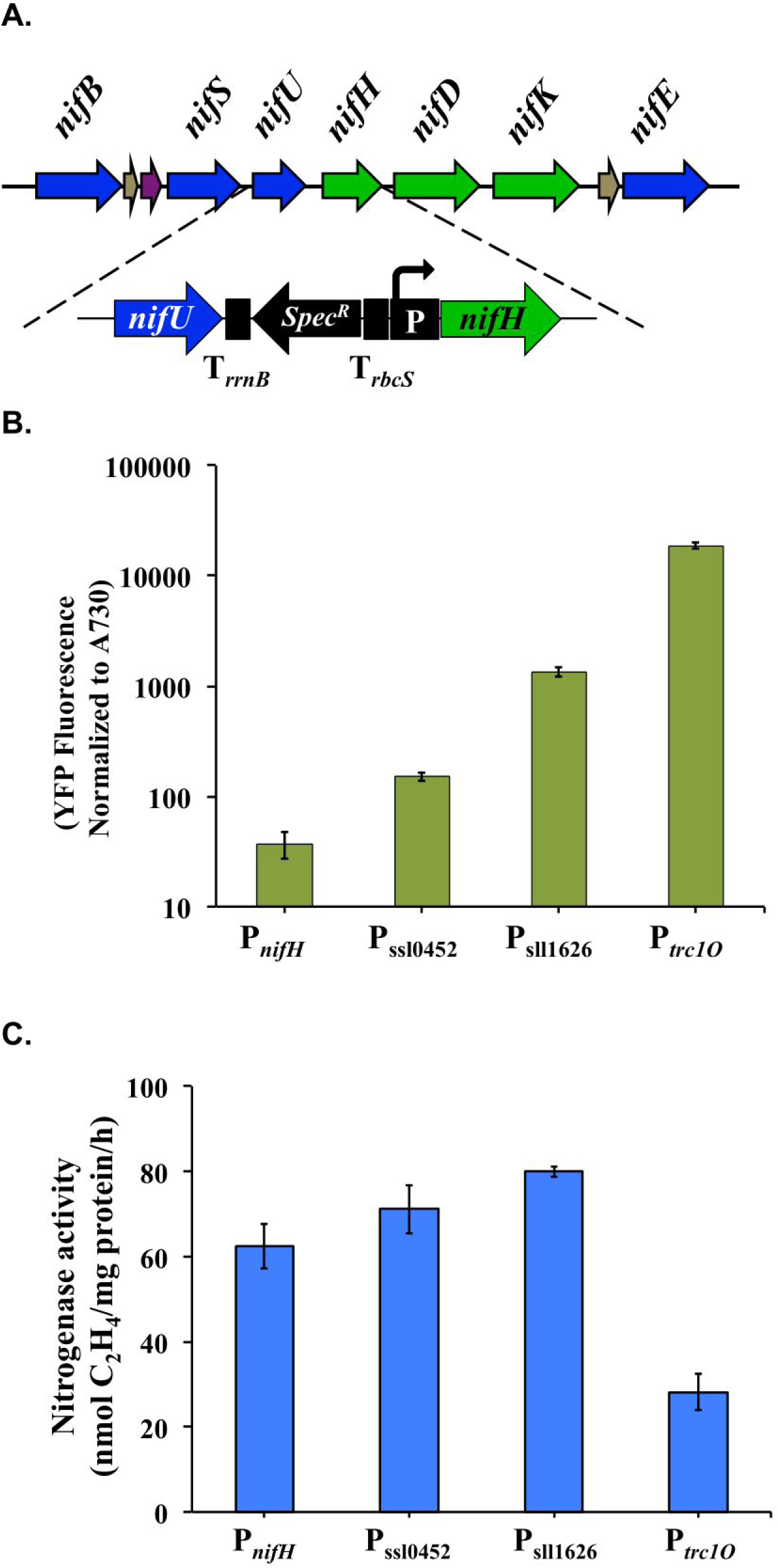
Expression levels of nitrogenase structural genes affect nitrogen fixation activities in engineered *Synechocystis* 6803. (A) Schematic map of changes on the 5’-UTR sequence before the *nifH* gene. Three promoters were tested, P_ssl0452_, P_sll1626_ and P*_trc1O_*. (B) The strengths of tested promoters were compared to the native promoter P*_nifH_*. The strengths were tested from the expression levels of the reporter protein EYFP. (C) Nitrogenase activity under different strength promoters preceding the *nifH* gene. Samples were collected from cultures under 12-h light/12-h dark conditions in BG11_0_ medium. Nitrogen fixation activity was assayed by acetylene reduction, and error bars represent the standard deviations observed from at least three independent experiments.

Three promoters characterized in an earlier study (17), P_sll0452_, P_sll1626_, and P*_trc1O_*, were tested for their efficacy in driving gene expression in comparison to the natural promoter P*_nifH_*. Using EYFP as the reporter protein (Fig. S1), we found that the strength of P*_nifH_* was the lowest among all the promoters tested (Fig. 1B). Nitrogenase activities could be enhanced when the stronger promoters P_sll0452_ and P_sll1626_ were used. However, when the strongest promoter P*_trc1O_* was used to drive the expression of *nifHDK*, more than 50% reduction in nitrogenase activity was observed (Fig. 1C). These results suggested that optimal expression of the structural *nif* genes is important for achieving high nitrogenase activity, a finding in accordance with a study in *Klebsiella* (20).

### Bottom-up strategy to re-organize nitrogen fixation genes

To find the essential genes for nitrogenase activity, we used a bottom-up strategy to build operons containing the genes from the native *Cyanothece* 51142 *nif* cluster (Fig. 2A). Promoters, RBS, and transcriptional terminators used in this study are native genetic elements of *Synechocystis* 6803 that were characterized earlier (17). The re-organized operons were inserted into the backbone of the shuttle vector pCB-SC101 (17).

**FIG 2.**
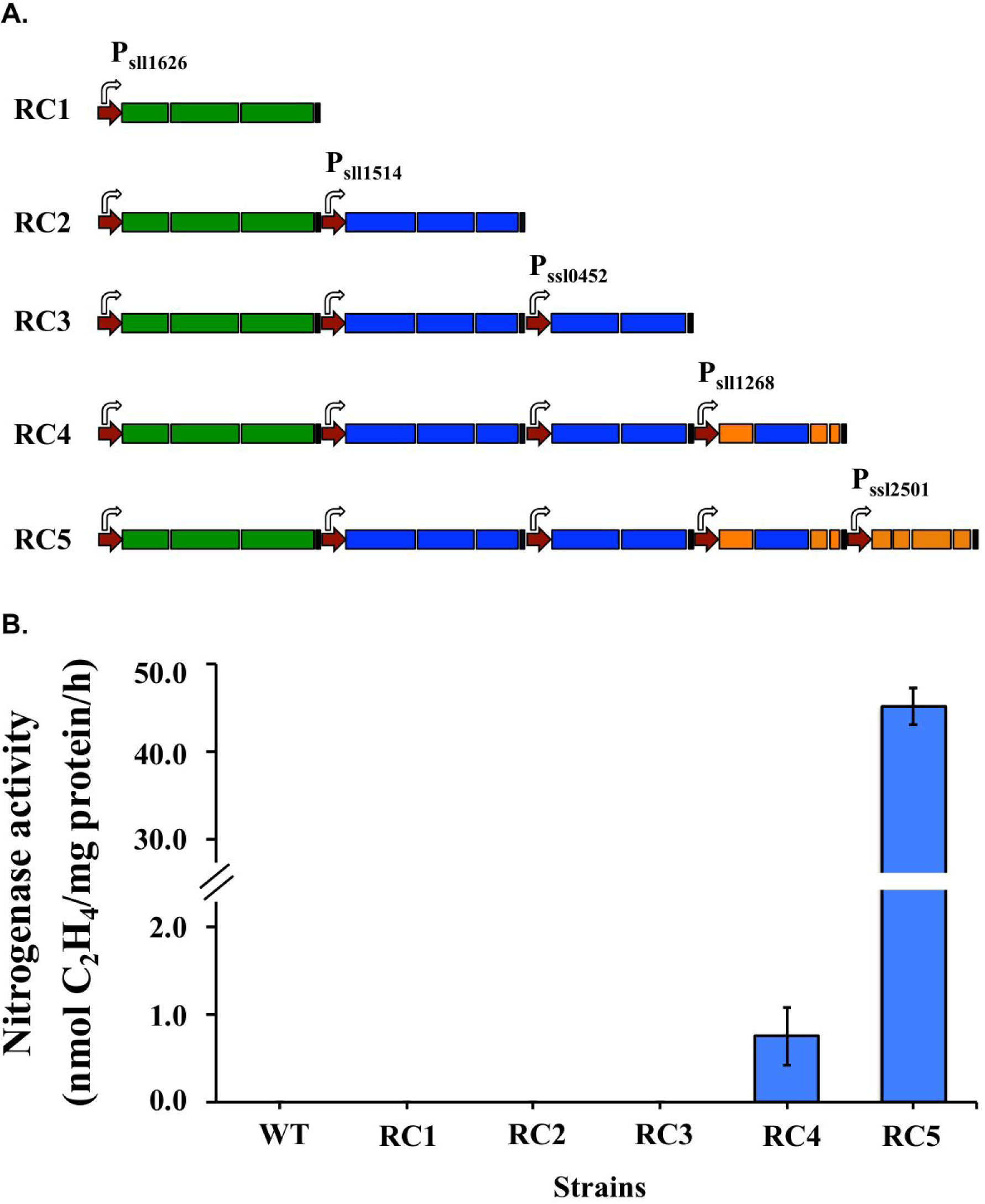
Rebuilding the functional nitrogenase enzyme in *Synechocystis* 6803. (A) A schematic map of re-construction of *nif* genes using a bottom-up strategy. Five operons were added sequentially into the plasmid pCB-SC101, which were transferred into *Synechocystis* 6803 as the CK strain. For each operon, promoters, RBS, and terminators were organized with ORFs to form an independent expression cassette. Shown are promoters (red), transcription terminators (black), and the genes for the three structural proteins, *nifHDK* (green), necessary cofactors (blue), accessory proteins (orange) (B) Nitrogenase activities of engineered strains containing sets of re-constructed *nif* operons. Samples were collected from cultures under 12-h light/12-h dark conditions in BG11_0_ medium. Nitrogen fixation activity was assayed by acetylene reduction, and error bars represent the standard deviations observed from at least three independent experiments.

Of the 24 genes in the native *Cyanothece nif* cluster, 16 genes have been annotated as *nif* genes in the JGI/IMG microbial database (<u>genome.jgi.doe.gov</u>) (21). The others are mostly genes of unknown function except one that is annotated as a ferredoxin gene. The 16 annotated genes were sequentially organized into 5 independent operons, *nifHDK*, *nifBSU*, *nifEN*, *nifPVZT*, and *nifXWhesAB* (Fig. S2). Five engineered strains (RC1 to RC5) were generated by transforming *Synechocystis* 6803 with plasmids containing the above operons in different combinations. The strain RC4 containing 12 genes (*nifHDKBSUENPVZT)* showed significant activity, generating 0.75 nmol ethylene per mg total protein in one hour (Fig. 2B). Complementing these 12 genes with the *nifXWhesAB* operon generated the RC5 strain, which exhibited a 60-fold increase in activity, thereby confirming these genes as essential components of the nitrogen fixing machinery. This observation was in accordance with our earlier findings (6) and the RC5 strain was selected for further analyses of the cluster.

### Influences of ferredoxins on nitrogenase activity

As direct electron donors to Fe protein, ferredoxins are known to significantly affect nitrogenase activity (22, 23). There is one ferredoxin, annotated as *fdxN*, in the 24-gene native *nif* cluster of *Cyanothece* 51142, and two more ferredoxins, *fdxH* and *fdxB*, in the 35-gene extended cluster. To evaluate the role of ferredoxins in nitrogen fixation, the three ferredoxin genes, *fdxN*, *fdxH*, and *fdxB*, were introduced into the RC5 strain. Four promoters, P_slr0701_, P_ssr2227_, P*_psbA2_*, and P*_trc1O_,* were used to independently drive the expression of each ferredoxin gene (Fig. 3A). One interesting observation was that, when driven by the P*_trc1O_* promoter, only the *fdxH* gene could be transformed into *Synechocystis*. Therefore, for genes *fdxN* and *fdxB*, only three promoters were tested.

**FIG 3.**
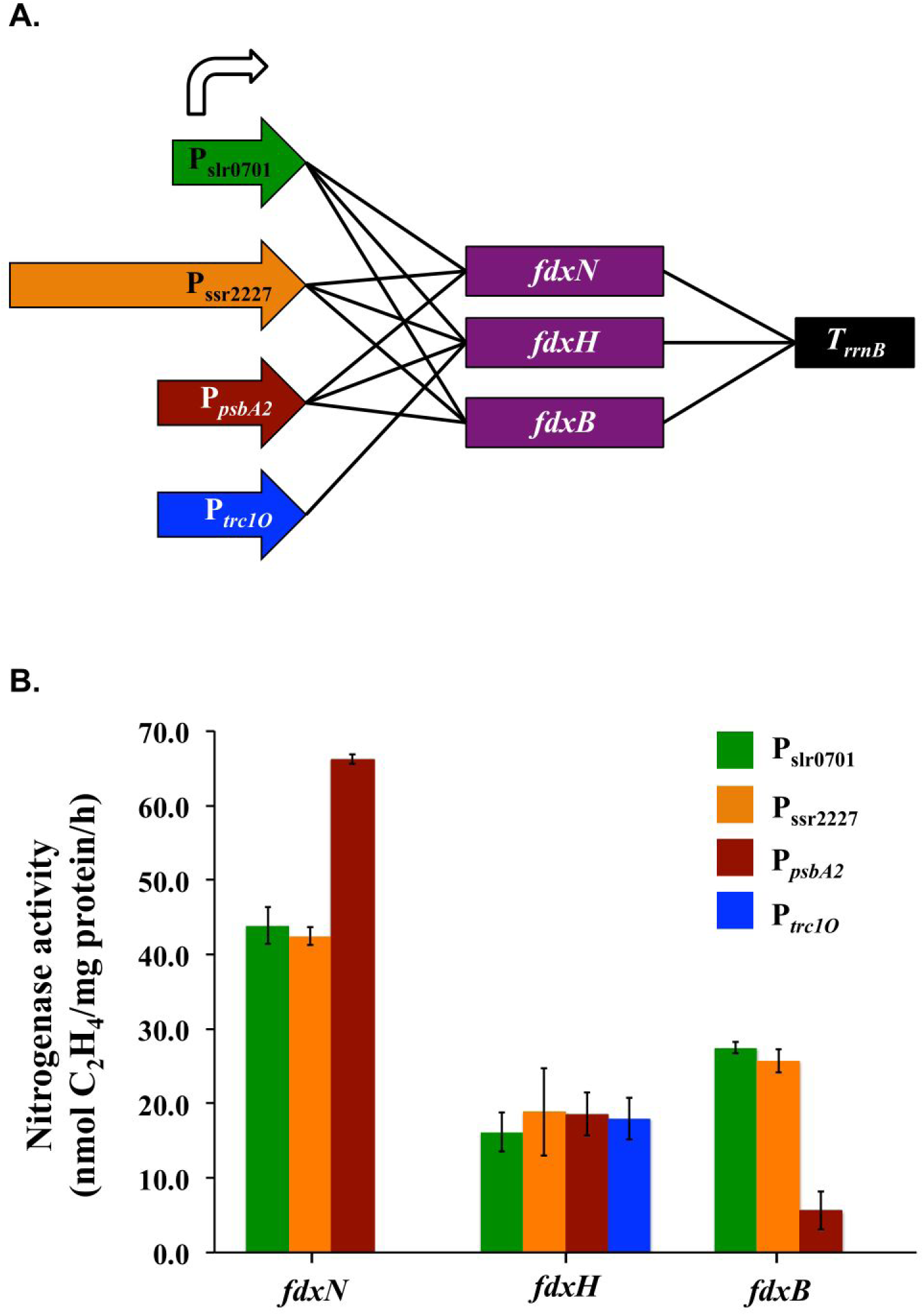
Ferredoxins affect nitrogenase activity in engineered *Synechocystis* 6803. (A) Scheme showing the combination of promoters and ferredoxin genes. The expression cassettes were integrated into the plasmid pRSF1010. (B) Nitrogenase activities are affected by type and expression level of ferredoxins in engineered *Synechocystis* 6803. Samples were collected from cultures under 12-h light/12-h dark conditions in BG11_0_ medium. Nitrogen fixation activity was assayed by acetylene reduction, and error bars represent the standard deviations observed from at least three independent experiments.

The ferredoxins tested had a diverse effect on nitrogenase activity (Fig. 3B). When the *fdxN* gene under the P*_psbA2_* promoter was introduced into RC5, a 50% increase in nitrogenase activity was observed. In contrast, when *fdxB* under a strong promoter was introduced into RC5, a significant reduction in activity was observed. Interestingly, introduction of *fdxH* into RC5 lead to a 60% decrease in activity, irrespective of the promoter used. This implied that the *fdxH* gene product negatively impacts nitrogen fixation.

### The essential *nif* genes for nitrogenase activity in *Synechocystis* 6803

Of the 16 annotated genes in strain RC5, only the structural genes *nifHDK* have been functionally characterized. The role of the other 13 genes in nitrogen fixation remains largely unknown. To determine the essential set of genes for nitrogenase activity in *Synechocystis* 6803, we used the CRISPR/cpf1 strategy (18) to seamlessly delete each of the 13 genes after the *nifHDK* operon (Fig. S3). The 13 mutant strains based on RC5 were generated as shown in Fig. 4A.

**FIG 4.**
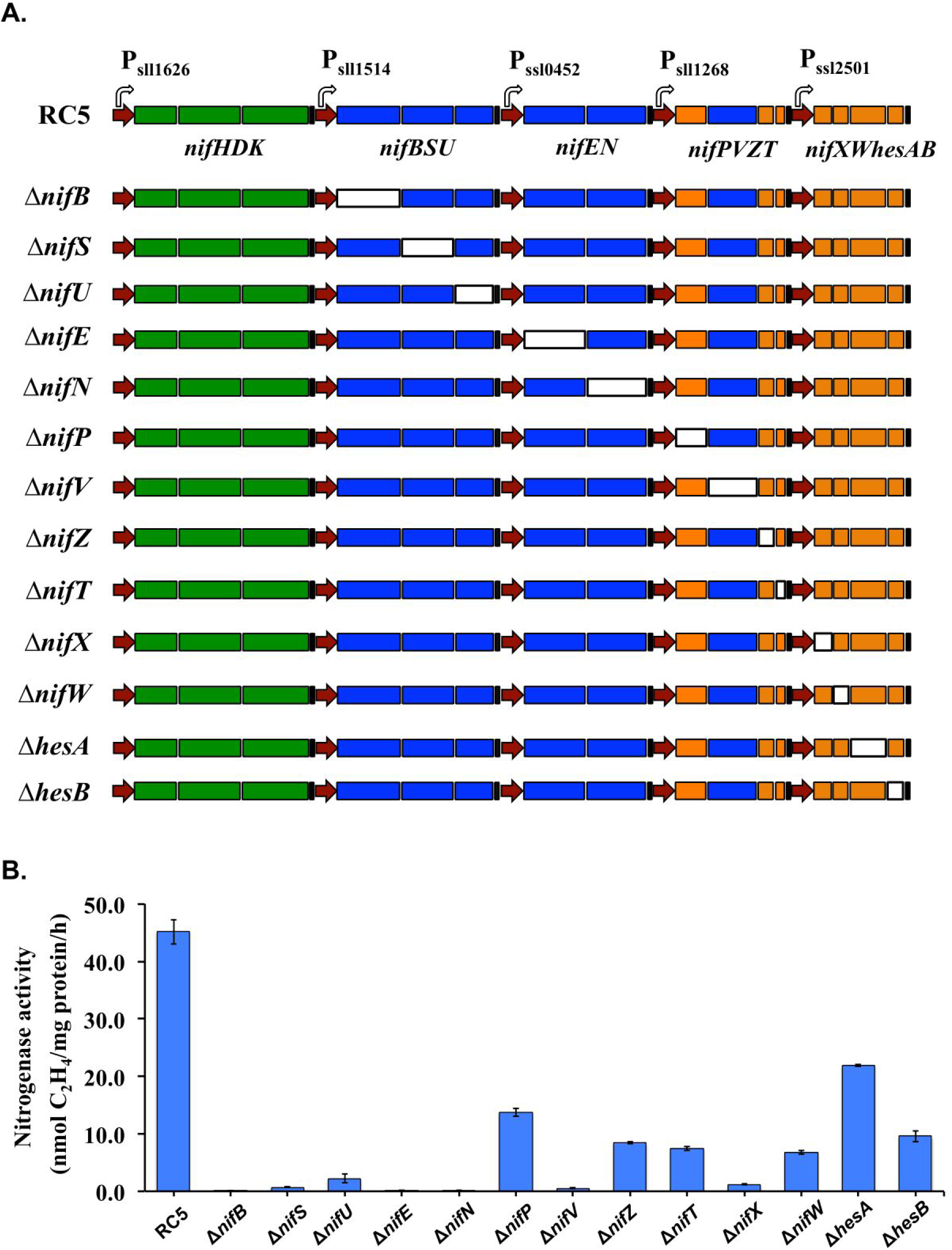
Deletion analysis of the genes shown in the re-constructed *nif* operons. (A) Scheme showing the deletion of specific genes in the operons, using the CRISPR/cpf1 system. The hollow rectangles represent the deleted genes, and the colored rectangles represent the remaining genes. (B) Nitrogenase activity of the deletion strains in engineered *Synechocystis* 6803. Samples were collected from cultures under 12-h light/12-h dark conditions in BG11_0_ medium. Nitrogen fixation activity was assayed by acetylene reduction, and error bars represent the standard deviations observed from at least three independent experiments.

As anticipated, deletion of *nifB*, *nifE*, or *nifN* lead to complete cessation of nitrogenase activity (Fig. 4B). This observation is in agreement with the fact that *nifENB* genes are required for M cluster biosynthesis and they are the most conserved genes besides *nifHDK* among all diazotrophs (11, 24). Similarly, deletion of any of the three genes *nifSU* and *nifV* resulted in over 100-fold decrease in nitrogenase activity. NifSU proteins are required for synthesis of iron-sulfur clusters (25), while *nifV* encodes the enzyme homocitrate synthase. Homocitrate is a component of the M cluster present in the catalytic center of dinitrogenase (26). NifP encodes a serine acetyltransferase, which has been identified as a key enzyme in cysteine biosynthesis in *E. coli* (27). Deletion of *nifP* (*cysE2*) resulted in approximately 60% reduction in nitrogenase activity in *Synechocystis*, implying its essential role in the process. Genes *nifZTW* encode proteins that are suggested to be involved in MoFe protein maturation, but none were found to be essential for nitrogen fixation in *E. coli* (13–15). However, our results showed that deletion of *nifZ,* lead to 70% decrease in nitrogenase activity, while deletion of *nifT* and W lead to about 80% decrease, implying their essential roles in nitrogen fixation in *a* photoautotroph. NifX is an interesting protein that has been identified as having various roles in different diazotrophs. It is a negative regulator of nitrogenase activity in *Klebsiella* (28), while in *Azotobacter* it is required for biosynthesis of the M cluster (29). Deletion of *nifX* in *Synechocystis* 6803 resulted in an over 100-fold decrease in nitrogenase activity, implying its essential role. Genes *hesAB* are not widely found in diazotrophs and their functions are yet to be determined. Our results suggested that they play an important role in nitrogenase activity in *Synechocystis* 6803; especially *hesB*, a finding corroborated by studies in the diazotrophic cyanobacterium *Anabaena* sp. PCC7120 (30).

## DISCUSSION

Members of the genus *Cyanothece* that engage in aerobic nitrogen fixation harbor one of the largest *nif* clusters known among cyanobacteria (31). The fact that these unicellular cyanobacteria are preferred by higher photoautotrophs as their nitrogen fixing partners suggests that their *nif* cluster might confer certain advantages. This hypothesis is further strengthened by recent studies that revealed that close relatives of *Cyanothece* with highly similar *nif* clusters are now functioning as early-stage nitrogen fixing organelles called diazoplasts or nitroplasts (1, 4, 32). Therefore, efforts to engineer photoautotrophic systems involving eukaryotic algae, lichens or plants for the production of nitrogen rich compounds will benefit from a detailed analysis of the *Cyanothece nif* cluster. In an earlier study we investigated the efficacy of the *Cyanothece* 51142 *nif* cluster in imparting nitrogen fixing ability to non-diazotrophic photoautotrophs by introducing it into *Synechocystis* 6803. The engineered *Synechocystis* strain exhibited significant rates of nitrogen fixation, laying the foundation for the current study, which further investigates the functioning of the cluster in photoautotrophs. We focused on identifying a set of essential genes and the optimal expression ratios of some of the key genes involved in the process of nitrogen fixation as the next step towards designing an efficient synthetic cluster for nitrogenase. We successfully re-organized the genes of this cluster to form independent operons in *Synechocystis* 6803 and demonstrated significant nitrogenase activity. Although the highest nitrogenase activity was observed in strain RC5, strain RC4 harboring only 12 *nif* genes also fixed nitrogen at low levels. In a study that introduced the *nif* gene cluster from *Leptolyngbya boryana* into *Synechocystis* 6803, nitrogenase activity around 4 nmol ethylene per mg cell dry weight per hour was demonstrated (33). We determined that under the testing conditions, the ratio of total protein per cell dry weight is 0.25 (data not shown), which translates into an activity of about 1 nmol ethylene per mg total protein in one hour, a rate very close to that observed in the RC4 strain. Although our engineered RC5 strain exhibits significantly lower activities (45 nmol per mg total protein in one hour) compared to *Cyanothece* 51142 (1262 nmol per mg total protein in one hour), the activities are significantly higher compared to the *Synechocystis* 6803 strain expressing genes from *Leptolyngbya boryana* (1 nmol per mg total protein in one hour) (33). Thus, the nitrogen fixation rates observed in the *Synechocystis* strain expressing genes from *Cyanothece* 51142 are very encouraging and validate our hypothesis that the *Cyanothece nif* cluster offers an optimal starting point for designing synthetic solar powered nitrogen fixing systems.

In order to convert N_2_ into ammonia, reducing power is required by the nitrogenase enzyme complex. Ferredoxins have been identified as one of the primary electron donors to the connected Fe protein of nitrogenase (34). However, in strain RC5, none of the ferredoxins from *Cyanothece* 51142 were transferred into *Synechocystis* 6803, which indicates that the endogenous ferredoxins in *Synechocystis* 6803 recognize the introduced nitrogenase Fe protein and function as the electrons bridge for dinitrogen gas reduction. It has not been determined which ferredoxin gene is essential for electron transfer to nitrogenase in *Cyanothece* 51142, and therefore all three annotated genes in the extended cluster were tested in the RC5 strain. Our results indicated that only the *fdxN* gene positively impacted nitrogenase activity when expressed at a higher level, suggesting that FdxN could be involved in electron transfer to nitrogenase. Studies have revealed that pathways of electron transfer to nitrogenase in metabolically diverse strains differ and the oxygen dynamics of the cell is an important factor that determines the ferredoxins involved in the process (35). In our study FdxH and FdxB negatively impacted nitrogenase activity, implying that these ferredoxins could be involved in diverting electrons from nitrogen fixation to other processes in *Synechocystis* 6803 where the cellular oxygen dynamics are very different from *Cyanothece* 51142. It has been shown that electron transport chains from plant organelles function to support nitrogenase activity in engineered *E. coli* (23). The fact that the RC5 strain fixes nitrogen without any ferredoxins from *Cyanothece* suggests that transferring *nif* genes without the ferredoxins from the native cluster might endow higher phototrophs with nitrogenase activity, minimizing the number of genes to be transferred.

In this work, we have described the minimal set of *nif* genes essential for nitrogenase activity in a nondiazotrophic cyanobacterium. However, the complete cellular machinery needed for robust and efficient nitrogen fixation under aerobic conditions, a key piece of the puzzle for engineering photoautotrophs, remains largely unknown. More recently, a transposon sequencing (Tn-Seq) strategy has been used in various non-diazotrophic species, including cyanobacteria, to identify essential gene sets under special conditions (36, 37). Extending such studies to diazotrophic cyanobacterial strains can reveal a more complete picture of the entire cellular machinery needed for high levels of nitrogen fixation.

## MATERIALS AND METHODS

### Microorganisms, culture conditions, and media

All cyanobacterial strains were cultured in 100-ml flasks with fresh BG11 medium and appropriate antibiotics (20 µg/ml kanamycin, 3 µg/ml gentamycin, or 20 µg/ml spectinomycin). Cells were pre-cultured at 30°C, under 50 µmol photons⋅ m^-2^⋅ s^-1^ constant light, with shaking at 150 rpm. For nitrogenase assays, pre-cultured cells were collected and washed with fresh BG11 medium without nitrate (BG11_0_) and resuspended in 500 ml fresh BG11_0_ medium. Cells were grown at 30°C with air bubbling under 12h light/dark conditions under 50 µmol photons⋅ m^-2^⋅ s^-1^ of light. All cloning was performed in *E*. *coli* strain XL1-Blue grown in LB medium in culture tubes or on agar plates at 37 °C, supplemented with 50 µg/ml kanamycin, 20 µg/ml gentamycin, or 30 µg/ml spectinomycin, as needed.

### Construction of recombinant plasmids and engineered strains

Plasmids and strains used in this study are listed in Table S1, and all primers are listed in Table S2. Plasmids were constructed by the Gibson isothermal DNA assembly method (38) using linear fragments purified from PCR products. Genomic DNA from the strain C*yanothece* 51142 and *Synechocystis* 6803 were used as templates for PCR. The genetic elements, including promoters, ribosome binding sites, and transcriptional terminators, were chosen from our previous study (17). The plasmid pUC118 (39) was used as the backbone to construct the plasmids used in the promoter assay for structural genes. The broad host plasmid pRSF1010 (40) was used as the backbone for the promoter - EYFP assays and for the evaluation of the ferredoxin coding genes. The shuttle vector described in our previous study, pCB-SC101, was used for the re-construction work using the bottom-up strategy (17). The plasmid pSL2680 constructed in our lab (18) was used for the knock-out experiments.

Plasmids based on pUC118 and pCB-SC101 vectors were all transformed by natural transformation (41) into the *Synechocystis* 6803 TSyNif-2 strain and wild-type strain, respectively. A tri-parental conjugation method was used to transfer all pRSF1010 derivative plasmids into *Synechocystis* 6803 cells (42) using a helper strain of *E. coli* containing the pRL443 and pRL623 plasmids (43). Transformants were selected on BG11 agar plates containing 20 µg/ml kanamycin, 3 µg/ml gentamycin, or 20 µg/ml spectinomycin, as needed. Isolated *Synechocystis* 6803 transformants were checked by PCR to confirm the presence of the desired constructs.

All PCR amplifications were performed using Phusion High-fidelity DNA polymerase (Thermo Scientifc). Plasmids and PCR products were purified using the GeneJET (Thermo Scientifc) plasmid miniprep kit and gel extraction kit, respectively. Oligonucleotides were designed using SnapGene software (GSL Biotech LLC) and synthesized by IDT (Coralville, IA). All of the plasmids constructed in this study were checked by sequencing to confirm the correct DNA coding information (Genewiz, NJ).

### Fluorescence measurements

Fluorescence intensities were measured as described earlier (17). Briefly, all cultures were adjusted to similar cell densities, with an OD_730 nm_ of 0.2 (about 1×10^8^ cells/ml) at the start of the experiment. Three independent replicates of each culture were then transferred to 6-well plates and grown for 3 days, followed by fluorescence measurements. The fluorescence intensity and the optical density of each culture were determined on a BioTek Synergy Mx plate reader (BioTek, Winooski, VT). The excitation and emission wavelengths were set to 485 and 528 nm for EYFP. All fluorescence data were normalized by culture density.

### Measurement of nitrogen fixation activity

Nitrogen fixation activity was measured by an acetylene reduction assay, modified from a previously published method (44). The assay was performed as follows: 25 ml of cyanobacterial culture grown in BG11_0_ medium was transferred to a 125 ml air-tight glass vial. 10 µM DCMU was added to the culture, vials were flushed with pure argon, and cultures were incubated in 12h light/dark conditions. Cells in the sealed vials were cultured overnight and at the time point D1, 5 ml acetylene was added into the sealed vials, followed by 3 hours of incubation in light at 30°C. 200 µl of gas was sampled from the headspace and injected into an Agilent 6890N Gas Chromatograph equipped with a Poropak N column and a flame ionization detector, using argon as the carrier gas. The temperature of the detector, injector, and oven were 200°C, 150°C and 100°C, respectively.

### Determination of cellular total proteins and cell dry weight

Total protein levels were determined on a plate reader (Bio-Tek Instruments, Winooski, VT) using a BCA-assay kit (Pierce, Rockford, IL) according to the manufacturer’s instructions. Cell dry weight (CDW) was determined from cell pellets of 100-ml culture aliquots that were centrifuged for 15 min at 4°C and 9,000×*g*, washed twice with distilled water, and dried at 85°C until the weight was constant.

## ACKNOWLEDGEMENTS

This study was supported by funding from the National Science Foundation (MCB-1331194) and by funding from the DOE Office of Science Energy Earthshot Initiative as part of the SFEE project under Award #DE-SC0024702. We thank Xiujun Duan for expert technical assistance and members of our laboratory for critical scientific discussions. D.L., M.B.P. and H.B.P. designed the experiments; D.L. performed the experiments; D.L., A.B., M.L., M.B.P. and H.B.P. wrote the paper.

## REFERENCES

1. Moulin SLY, Frail S, Braukmann T, Doenier J, Steele-Ogus M, Marks JC, Mills MM, Yeh E. 2024. The endosymbiont of *Epithemia clementina* is specialized for nitrogen fixation within a photosynthetic eukaryote. ISME Commun 4.

2. Prechtl J, Kneip C, Lockhart P, Wenderoth K, Maier U-G. 2004. Intracellular spheroid bodies of *Rhopalodia gibba* have nitrogen-fixing apparatus of cyanobacterial origin. Mol Biol Evol 21:1477–1481.

3. Carpenter EJ, Janson S. 2000. Intracellular cyanobacterial symbionts in the marine diatom *Climacodium Frauenfeldianum* (Bacillariophyceae). J Phycol 36:540–544.

4. Coale TH, Loconte V, Turk-Kubo KA, Vanslembrouck B, Mak WKE, Cheung S, Ekman A, Chen J-H, Hagino K, Takano Y, Nishimura T, Adachi M, Le Gros M, Larabell C, Zehr JP. 2024. Nitrogen-fixing organelle in a marine alga. Science 384:217–222.

5. Hagino K, Onuma R, Kawachi M, Horiguchi T. 2013. Discovery of an endosymbiotic nitrogen-fixing cyanobacterium UCYN-A in *Braarudosphaera bigelowii* (Prymnesiophyceae). PLoS One 8:e81749.

6. Liu D, Liberton M, Yu J, Pakrasi HB, Bhattacharyya-Pakrasi M. 2018. Engineering nitrogen fixation activity in an oxygenic phototroph. MBio 9.

7. Tezcan FA, Kaiser JT, Mustafi D, Walton MY, Howard JB, Rees DC. 2005. Nitrogenase complexes: multiple docking sites for a nucleotide switch protein. Science 309:1377–80.

8. Hu Y, Ribbe MW. 2013. Nitrogenase assembly. Biochimica et Biophysica Acta (BBA) 1827:1112–22.

9. Sickerman NS, Ribbe MW, Hu Y. 2017. Nitrogenase cofactor assembly: an elemental inventory. Acc Chem Res 50:2834–2841.

10. Sickerman NS, Rettberg LA, Lee CC, Hu Y, Ribbe MW. 2017. Cluster assembly in nitrogenase. Essays Biochem 61:271–279.

11. Dos Santos PC, Fang Z, Mason SW, Setubal JC, Dixon R. 2012. Distribution of nitrogen fixation and nitrogenase-like sequences amongst microbial genomes. BMC Genomics 13:162.

12. Esteves-Ferreira AA, Cavalcanti JHF, Vaz M, Alvarenga LV, Nunes-Nesi A, Araujo WL. 2017. Cyanobacterial nitrogenases: phylogenetic diversity, regulation and functional predictions. Genet Mol Biol 40:261–275.

13. Wang L, Zhang L, Liu Z, Zhao D, Liu X, Zhang B, Xie J, Hong Y, Li P, Chen S, Dixon R, Li J. 2013. A minimal nitrogen fixation gene cluster from *Paenibacillus* sp. WLY78 enables expression of active nitrogenase in *Escherichia coli*. PLoS Genet 9:e1003865.

14. Yang J, Xie X, Wang X, Dixon R, Wang YP. 2014. Reconstruction and minimal gene requirements for the alternative iron-only nitrogenase in *Escherichia coli*. Proc Natl Acad Sci U S A 111:E3718–25.

15. Yang J, Xie X, Xiang N, Tian ZX, Dixon R, Wang YP. 2018. Polyprotein strategy for stoichiometric assembly of nitrogen fixation components for synthetic biology. Proc Natl Acad Sci U S A 115:E8509–E8517.

16. Ito Y, Yoshidome D, Hidaka M, Araki Y, Ito K, Kosono S, Nishiyama M. 2024. Improvement of the nitrogenase activity in *Escherichia coli* that expresses the nitrogen fixation-related genes from *Azotobacter vinelandii*. Biochem Biophys Res Commun 728:150345.

17. Liu D, Pakrasi HB. 2018. Exploring native genetic elements as plug-in tools for synthetic biology in the cyanobacterium *Synechocystis* sp. PCC 6803. Microb Cell Fact 17:48.

18. Ungerer J, Pakrasi HB. 2016. Cpf1 Is a versatile tool for CRISPR genome editing across diverse species of cyanobacteria. Sci Rep 6:39681.

19. Colon-Lopez MS, Tang H, Tucker DL, Sherman LA. 1999. Analysis of the *nifHDK* operon and structure of the NifH protein from the unicellular, diazotrophic cyanobacterium, *Cyanothece* strain sp. ATCC 51142. Biochimica et Biophysica Acta (BBA) 1473:363-75.

20. Temme K, Zhao D, Voigt CA. 2012. Refactoring the nitrogen fixation gene cluster from *Klebsiella oxytoca*. Proc Natl Acad Sci U S A 109:7085–90.

21. Nordberg H, Cantor M, Dusheyko S, Hua S, Poliakov A, Shabalov I, Smirnova T, Grigoriev IV, Dubchak I. 2014. The genome portal of the Department of Energy Joint Genome Institute: 2014 updates. Nucleic Acids Res 42:D26–31.

22. Duval S, Danyal K, Shaw S, Lytle AK, Dean DR, Hoffman BM, Antony E, Seefeldt LC. 2013. Electron transfer precedes ATP hydrolysis during nitrogenase catalysis. Proc Natl Acad Sci U S A 110:16414–9.

23. Yang J, Xie X, Yang M, Dixon R, Wang YP. 2017. Modular electron- transport chains from eukaryotic organelles function to support nitrogenase activity. Proc Natl Acad Sci U S A 114:E2460–E2465.

24. Curatti L, Hernandez JA, Igarashi RY, Soboh B, Zhao D, Rubio LM. 2007. In vitro synthesis of the iron-molybdenum cofactor of nitrogenase from iron, sulfur, molybdenum, and homocitrate using purified proteins. Proc Natl Acad Sci U S A 104:17626–31.

25. Zhao D, Curatti L, Rubio LM. 2007. Evidence for *nifU* and *nifS* participation in the biosynthesis of the iron-molybdenum cofactor of nitrogenase. J Biol Chem 282:37016–25.

26. Rubio LM, Ludden PW. 2008. Biosynthesis of the iron-molybdenum cofactor of nitrogenase. Annu Rev Microbiol 62:93–111.

27. Denk D, Bock A. 1987. L-cysteine biosynthesis in *Escherichia coli*: nucleotide sequence and expression of the serine acetyltransferase (*cysE*) gene from the wild-type and a cysteine-excreting mutant. J Gen Microbiol 133:515–25.

28. Gosink MM, Franklin NM, Roberts GP. 1990. The product of the *Klebsiella pneumoniae nifX* gene is a negative regulator of the nitrogen fixation (*nif*) regulon. J Bacteriol 172:1441–7.

29. Shah VK, Rangaraj P, Chatterjee R, Allen RM, Roll JT, Roberts GP, Ludden PW. 1999. Requirement of NifX and other *nif* proteins for in vitro biosynthesis of the iron-molybdenum cofactor of nitrogenase. J Bacteriol 181:2797–801.

30. Borthakur D, Basche M, Buikema WJ, Borthakur PB, Haselkorn R. 1990. Expression, nucleotide sequence and mutational analysis of two open reading frames in the *nif* gene region of *Anabaena* sp. strain PCC7120. Mol Genet Genomics 221:227–34.

31. Welsh EA, Liberton M, Stockel J, Loh T, Elvitigala T, Wang C, Wollam A, Fulton RS, Clifton SW, Jacobs JM, Aurora R, Ghosh BK, Sherman LA, Smith RD, Wilson RK, Pakrasi HB. 2008. The genome of *Cyanothece* 51142, a unicellular diazotrophic cyanobacterium important in the marine nitrogen cycle. Proc Natl Acad Sci U S A 105:15094–9.

32. Bandyopadhyay A, Elvitigala T, Welsh E, Stockel J, Liberton M, Min H, Sherman LA, Pakrasi HB. 2011. Novel metabolic attributes of the genus *Cyanothece*, comprising a group of unicellular nitrogen-fixing *Cyanothece*. MBio 2.

33. Tsujimoto R, Kotani H, Yokomizo K, Yamakawa H, Nonaka A, Fujita Y. 2018. Functional expression of an oxygen-labile nitrogenase in an oxygenic photosynthetic organism. Sci Rep 8:7380.

34. Poudel S, Colman DR, Fixen KR, Ledbetter RN, Zheng Y, Pence N, Seefeldt LC, Peters JW, Harwood CS, Boyd ES. 2018. Electron transfer to nitrogenase in different genomic and metabolic backgrounds. J Bacteriol doi:10.1128/jb.00757-17.

35. Poudel S, Colman DR, Fixen KR, Ledbetter RN, Zheng Y, Pence N, Seefeldt LC, Peters JW, Harwood CS, Boyd ES. 2018. Electron transfer to nitrogenase in different genomic and metabolic backgrounds. J Bacteriol 200.

36. Luo H, Lin Y, Gao F, Zhang CT, Zhang R. 2014. DEG 10, an update of the database of essential genes that includes both protein-coding genes and noncoding genomic elements. Nucleic Acids Res 42:D574–80.

37. Rubin BE, Wetmore KM, Price MN, Diamond S, Shultzaberger RK, Lowe LC, Curtin G, Arkin AP, Deutschbauer A, Golden SS. 2015. The essential gene set of a photosynthetic organism. Proc Natl Acad Sci U S A 112:E6634–43.

38. Gibson DG, Young L, Chuang RY, Venter JC, Hutchison CA, 3rd, Smith HO. 2009. Enzymatic assembly of DNA molecules up to several hundred kilobases. Nat Methods 6:343-5.

39. Vieira J, Messing J. 1987. Production of single-stranded plasmid DNA. Methods Enzymol 153:3–11.

40. Taton A, Unglaub F, Wright NE, Zeng WY, Paz-Yepes J, Brahamsha B, Palenik B, Peterson TC, Haerizadeh F, Golden SS, Golden JW. 2014. Broad-host-range vector system for synthetic biology and biotechnology in cyanobacteria. Nucleic Acids Res 42:e136.

41. Williams JGK. 1988. Construction of specific mutations in photosystem II photosynthetic reaction center by genetic engineering methods in Synechocystis 6803, p 766-778, Methods Enzymol, vol Volume 167. Academic Press.

42. Golden SS, Brusslan J, Haselkorn R. 1987. Genetic engineering of the cyanobacterial chromosome. Methods Enzymol 153:215–31.

43. Tsinoremas NF, Kutach AK, Strayer CA, Golden SS. 1994. Efficient gene transfer in *Synechococcus* sp. strains PCC 7942 and PCC 6301 by interspecies conjugation and chromosomal recombination. J Bacteriol 176:6764–8.

44. Bandyopadhyay A, Stockel J, Min H, Sherman LA, Pakrasi HB. 2010. High rates of photobiological H_2_ production by a cyanobacterium under aerobic conditions. Nat Commun 1:139.

